# Amygdalostriatal coupling underpins positive but not negative coloring of ambiguous affect

**DOI:** 10.1101/811505

**Authors:** M. Justin Kim, Alison M. Mattek, Jin Shin

**Author notes:** Corresponding Author: M. Justin Kim, Ph.D., Department of Psychology, Sungkyunkwan University, Seoul, South Korea 03063.

## Abstract

Humans routinely integrate affective information from multiple sources. For example, we rarely interpret an emotional facial expression devoid of context. Here, we describe the neural correlates of an affective computation that involves integrating multiple sources, by leveraging the ambiguity and subtle feature-based valence signals found in surprised faces. Using functional magnetic resonance imaging, participants reported the valence of surprised faces modulated by positive or negative sentences. Amygdala activity corresponded to the valence value assigned to each contextually modulated face, with greater activity reflecting more negative ratings. Amygdala activity did not track the valence of the faces or sentences *per se*. Moreover, the amygdala was functionally coupled with the nucleus accumbens only during face trials preceded by positive contextual cues. These data suggest 1) valence-related amygdala activity reflects the integrated valence values rather than the valence values of each individual component, and 2) amygdalostriatal coupling underpins positive but not negative coloring of ambiguous affect.

## Introduction

Affective valence – the degree of positivity-negativity or pleasantness-unpleasantness – is one of the core dimensions that underpins any emotional stimulus, event, or experience (Russell, 1980; Cacioppo et al., 1986; Lang, Bradley, & Cuthbert, 1998). Neural encoding of valence is traditionally known to occur in several brain regions centered on the limbic system and its interconnected network of cortical areas, which includes the amygdala-prefrontal circuitry. For example, single-cell recording studies of nonhuman primates showed that neurons in the amygdala (Belova, Paton, & Salzman, 2008; Paton et al., 2006), as well as the orbitofrontal cortex (OFC) (Morrison & Salzman, 2009) encode negative and positive valence. Human neuroimaging research also found that activity of the amygdala (Anders et al., 2008; Comte et al., 2016; Costafreda et al., 2008; Cunningham, Van Bavel, & Johnsen., 2008; Jin et al., 2015; Kim et al., 2017; Phelps & LeDoux, 2005), ventromedial prefrontal cortex (vmPFC)/OFC (Chikazoe et al., 2014; Clithero & Rangel, 2013; Gottfried, O’Doherty, & Dolan, 2002) and the dorsomedial prefrontal cortex (dmPFC) (Etkin, Egner, & Kalisch., 2011; Skerry & Saxe, 2014) was associated with the valence value of emotional stimuli. Furthermore, studies utilizing multivoxel pattern analysis (MVPA) of functional magnetic resonance imaging (fMRI) data and meta-analyses of human neuroimaging studies showed that brain regions beyond the corticolimbic areas, such as the precuneus, superior temporal sulcus, thalamus, anterior insula, and the sensory cortex may also play a role in encoding modality-general or modality-specific valence values (Chang et al., 2015; Kim, Shinkareva, & Wedell, 2017; Knutson, Katovich, & Suri, 2014; Lindquist et al., 2016; Mattek et al., 2020; Miskovic & Anderson 2018).

In our previous work we demonstrated that the amygdala was able to track the valence signal embedded within facial features, and the use of emotionally ambiguous surprised faces guaranteed this effect was unconfounded by the arousal dimension or emotion category (Kim et al., 2017; Mattek, Wolford, & Whalen, 2017). Though this study and the aforementioned work have collectively further elucidated the link between valence and patterns of neural activity, they have primarily focused on responses to a single source of emotion (single stimulus presentations of photos of emotion-eliciting scenes, facial expressions of emotion, or pleasant/unpleasant odor, etc.). It remains to be seen how valence is represented in the brain when it is computed by integrating multiple sources of affective information. To put it another way, it is possible that the increased amygdala response to negative valence values observed in Kim et al. (2017) reflects either 1) valence signals related to low-level features (i.e., facial features) only, or 2) the subjective affective evaluation or judgment of those features. The critical difference between the two accounts is that the former is independent of the availability of other affective information, whereas the latter relies on them. In the case of Kim et al. (2017) and many other studies that relied on a single source of affective information to extract valence values, these two accounts of amygdala function are indistinguishable, as both would equally explain the observed association between valence and amygdala activity. Here, we use a task that involves multiple sources of affective information to achieve meaningful delineation of the two possibilities.

To achieve this, we adopted a paradigm from a previous study (Kim et al., 2004), which showed that clearly-valenced contextual information shifts the perception of an ambiguously-valenced low-level signal (e.g., providing a scenario that depicts a negative situation such as “He lost $500” paired with a surprised face). The perceptual shift in the perceived valence of a contextually-modulated surprised face in such case would be a result of a psychological process, or *valence computation*, in which affective information from contextual cues and facial features are integrated. It has been shown that surprised faces modulated with negative compared to positive verbal context elicit increased amygdala activity (Kim et al., 2004). Such behavior of the amygdala is generalizable to nonverbal contextual cues (e.g., negative pictures, fearful faces) coloring the affective perception of neutral faces (Grupe et al., 2018; Lapate et al., 2016; McRae et al., 2012). Based on this, we sought to test the possibility that the amygdala, at least at the level of activity that can be measured with fMRI, may reflect the final, subjective judgment of valence, rather than the valence value of context or features alone. Furthermore, there may be observable differences in the functional connectivity patterns of the amygdala depending on the valence of a given context, which could shed light onto a possible neural mechanism that underpins the integration of affective information from top-down (e.g., context) and bottom-up (e.g., facial features) sources.

Of particular neuroanatomical interest to the latter point is the ventral striatum. Ventral areas of the striatum include the nucleus accumbens (NAcc), which is well known for its role in positive affect, desire, reward, and valuation (Berridge & Kringelbach, 2015; Wang, Smith, & Delgado, 2016; see also Reynolds & Berridge, 2008 for NAcc’s role in oppositely-valenced aversive behavior). The NAcc has extensive connections with the amygdala, which are mostly characterized as afferent inputs from the amygdala’s basolateral nucleus (Alheid & Heimer, 1988; Pitkänen, 2000; Rigoard et al., 2011). Animal studies have consistently demonstrated the functional importance of this amygdalostriatal projection in processing positive valence (Beyeler et al., 2018; Namburi et al., 2015; Stuber et al., 2011). Based on these findings, we sought to test the idea that the NAcc and the amygdala may actively communicate via an amygdalostriatal pathway during valence computation, particularly when affective information from facial features become integrated with positive contextual information.

Here, we used fMRI to test whether the activity of the amygdala encoded the combined valence of contextually modulated surprised faces, and/or the valence of the components - sentences and facial features. If amygdala activity reflects the subjective evaluation or judgment with respect to valence, then the data would support the former prediction. Conversely, if the amygdala primarily encodes the valence values that are generated from the bottom-up, then the data would follow the latter prediction, tracking the valence values embedded within the facial features. In both cases, we predicted that the amygdala would show increased activity to negative valence, based on previous fMRI studies reporting this effect unconfounded by arousal (Anders et al., 2008; Kim et al., 2004; Mattek et al., 2020). Based on the idea that the information from context and features are integrated during valence computation, we also examined whether the amygdala would show increased functional coupling with the NAcc when incorporating positive, but not negative contextual information. The main hypotheses were that 1) valence computed from multiple sources of affective information would be reflected in the amygdala, with more negative valence values corresponding to monotonic increase in activity; and 2) the amygdala would show greater functional connectivity with the NAcc only when the faces were modulated by positive contextual cues.

## Method

### Participants

Twenty-seven undergraduate students participated in the current study. Data from three participants were removed from further analyses due to excessive head movement during the fMRI scanning sessions (> 1.5 mm). Thus, a total of 24 healthy volunteers (15 females, ages 18-21 years, mean age = 19 years) were included in the current study. Given the within-subjects design employed in the present study, this sample size (*n* = 24) corresponded to the number of participants needed to attain 80% power to detect amygdala findings, based on a power analysis using a standard effect size that is associated with typical emotional tasks (Cohen’s *d* = ∼0.6) (Poldrack et al., 2017). All participants were screened for past or current psychiatric illnesses (Axis I or II) using the Structured Clinical Interview for DSM-IV (SCID) (First et al., 1995). No participants had any history of taking psychotropic medications. Prior to the experiment, all participants gave written, informed consent in accordance with the guidelines set by the Committee for the Protection of Human Subjects at Dartmouth College.

### Stimuli

A total of 36 surprised faces were selected from the Pictures of Facial Affect (Ekman & Friesen, 1976), NimStim (Tottenham et al., 2009), and Karolinska Directed Emotional Faces (Lundqvist, Flykt, & Ohman, 1998) facial expression sets, which were identical to those used in our previous fMRI study (Kim et al., 2017). These 36 identities were categorized as having *positive features (consensus positive), ambiguous features (no consensus)*, or *negative features (consensus negative)*. These three categories (12 identities in each), which were defined independently from our previous study (Kim et al., 2017), were based on the ratio of negative ratings each face received from a two-alternative forced choice task. These labels reflect the varying degree of valence signals embedded within the facial features of surprise (Kim et al., 2017). Previous work has shown that categorical classifications of valence can predict over 95% of the variance in continuous valence values (Mattek, Wolford, & Whalen, 2017). The rationale for choosing these 36 identities was twofold: 1) to facilitate comparison with our previous fMRI study, in which only the faces were presented without contextual cues, and 2) to explore the potential effects of feature-based valence of the faces *per se* and their interaction with contextual cues. A total of 12 sentences (6 positive, 6 negative), which were matched for absolute valence and arousal, were adopted from a previous study (Kim et al., 2004). Full list of the sentences is available in the appendix of Kim et al. (2004).

### Experimental Procedure

A modified version of a context modulation paradigm from Kim et al. (2004) was used for the current study. Modification included introducing 1) jittered inter-stimulus interval (2.5-3.6 s) between the presentation of a sentence and a face, 2) a response period during which participants rate the preceding face using a 9-point Likert rating scale, and 3) jittered inter-trial interval (2.5-3.6 s) between the rating trial and the appearance of the next sentence, to allow for analyses of rapid event-related fMRI data (Figure 1). We note that these jittered inter-stimulus interval and inter-trial interval timings were sufficient in mitigating collinearity across the sentence and face trials (all cos(*θ*) < 0.28; mean VIF = 1.36). Using E-Prime software, all of the stimuli were back projected onto a screen, on which the participants viewed during fMRI scanning through a mirror that was mounted on the head coil. All faces were gray scaled and normalized for size and luminance.

**Figure 1.**
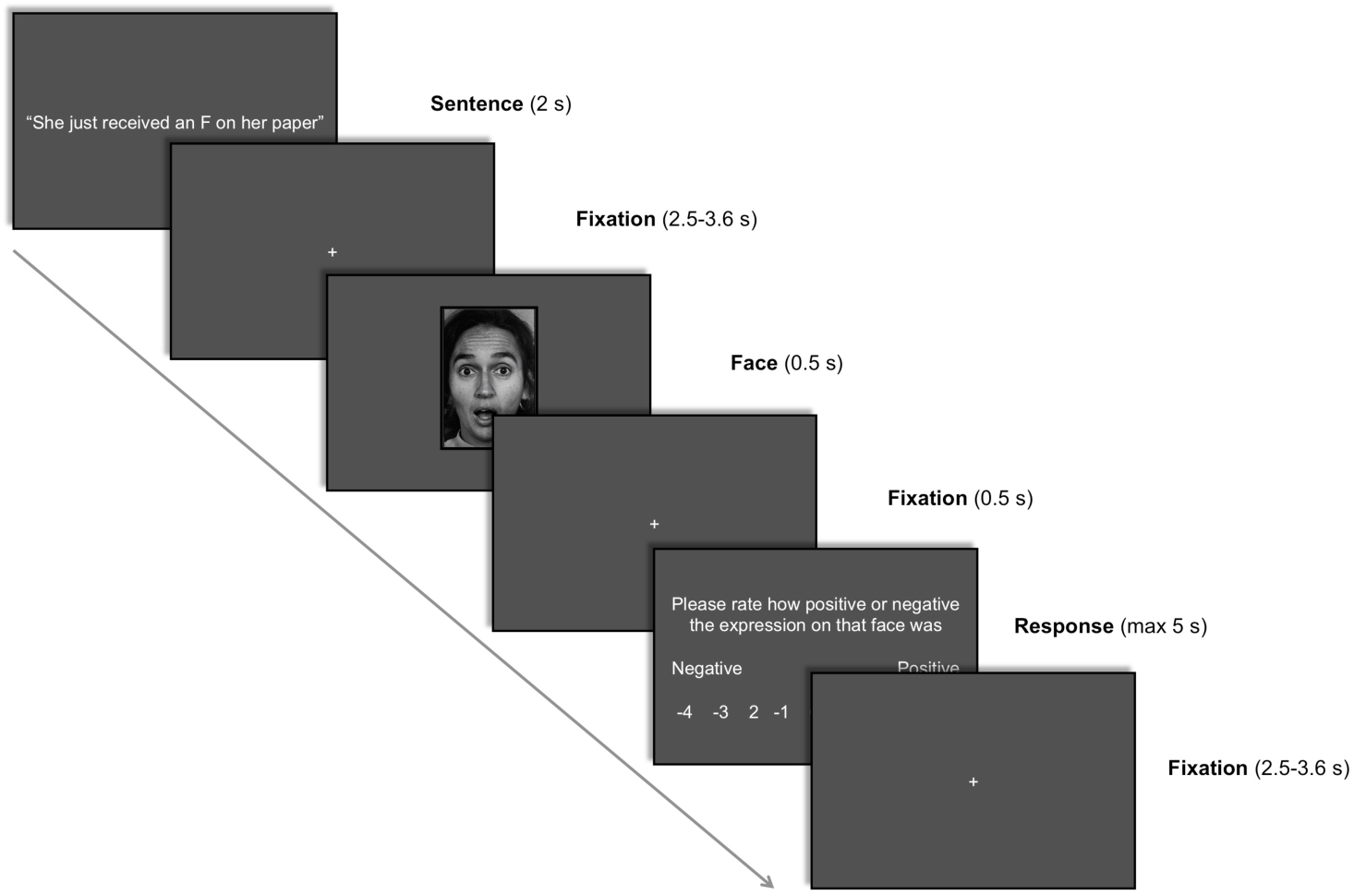
An example trial structure used in the fMRI experiment. Participants were told that the sentence describes a short scenario explaining why the person is making that expression. Then, they were carefully instructed to rate how positive or negative the expression on that face was.

During fMRI scanning, participants viewed pairs of sentences and surprised faces. The sentences provided verbal context by briefly explaining why the following face is making a surprised expression. Each sentence had two versions – positive and negative (e.g., “He just found $500” and “He just lost $500”), which were matched for absolute valence and arousal ratings (Kim et al., 2004). The sentences were always presented before the faces, which were followed by a 9-point Likert rating scale (ranging from -4 to +4; -4 = *very negative*, +4 = *very positive*) during which the participants were asked to report the valence of the *surprised face* using a MR compatible response box. Using two buttons, participants were able to select their responses on each trial by moving the slider in either direction of the scale. Once the participants selected the desired response, they were instructed to wait for 2 seconds before the next trial. The rating scale remained on the screen for a maximum of 5 seconds; when the response was made (i.e., remaining on a desired selection for 2 seconds), the trial then ended and then proceeded to the next trial. Sentences and faces were presented on the screen for 2,000 ms and 500 ms, respectively. Each of the 36 surprised faces was paired with two positive and two negative versions of the sentence, yielding a total of 144 trials, which was evenly split into four 8-minute runs. In sum, there were a total of six conditions: 1) positive features + positive context, 2) positive features + negative context, 3) ambiguous features + positive context, 4) ambiguous features + negative context, 5) negative features + positive context, 6) negative features + negative context.

### Image Aquisition

All participants were scanned at the Dartmouth Brain Imaging Center using a 3.0 Tesla Philips Intera Achieva Scanner (Philips Medical Systems, Bothell, WA) equipped with a 32-channel head coil. High-resolution anatomical T1-weighted magnetic resonance images were scanned using a magnetization-prepared rapid gradient echo sequence (MP-RAGE), with 220 sagittal slices (TE = 3.7 ms, TR = 8.2 ms, FOV = 240 mm, flip angle = 8**°**, voxel size = 0.9375 × 0.9375 × 1 mm). Functional images were acquired using echo-planar T2*-weighted imaging sequence. Each volume consisted of 33 interleaved 3-mm thick slices with 0.5 mm interslice gap (TE=35 ms, TR=2,000 ms, FOV=240 mm, flip angle=90°, voxel size=3×3×3.5 mm).

### fMRI Data Analysis

All fMRI blood-oxygen-level-dependent (BOLD) data were first preprocessed using SPM8 (Wellcome Department of Imaging Neuroscience, London, UK). Raw functional images were corrected for slice timing and head movement. None of the remaining 24 participants had head movement more than 1.5 mm in any direction. Functional images were then normalized to standard space (3×3×3 mm) using the Montreal Neurological Institute (MNI)-152 template. Spatial smoothing was applied to the normalized functional images using a Gaussian kernel of 6 mm full width at half maximum.

For each participant, two separate general linear models (GLM) were constructed: 1) a GLM that incorporated a parametric modulator (idiosyncratic valence ratings for each trial) of the height of the hemodynamic response during the face trials (GLM_1_), and 2) a GLM that incorporated the six task conditions (modeled at the onset of the faces) plus the two context types (modeled at the onset of the sentences), convolved with a hemodynamic response function (GLM_2_). GLM_1_ utilized a parametric modulation analysis approach to assess whether BOLD responses were increased or decreased as a function of trial-by-trial valence ratings, whereas GLM_2_ tests for overall differences across conditions, not taking into account the trial-by-trial valence ratings. Covariates of no interest (a session mean, a linear trend for each run to account for low-frequency drift, and six movement parameters derived from realignment corrections) were also included in both GLMs. Linear contrast maps for the parametric modulator (GLM_1_), each condition versus fixation (GLM_2_), as well as negatively modulated surprise versus positively modulated surprise (i.e., contextual modulation effects; GLM_2_) and surprise faces with negative features versus positive features (i.e., feature-based valence effects; GLM_2_) were generated for each participant. The latter two contrasts were selected *a priori* to facilitate direct comparison with the findings from Kim et al. (2004) and Kim et al., (2017), respectively, in which the same contrasts were used to observe changes in amygdala activity. Contrast maps were then entered into a random effects model, which accounts for inter-subject variability and allows population-based inferences to be drawn. In addition, to accommodate the 2 × 3 factorial design of the current experiment, a voxelwise 2 × 3 analysis of variance (ANOVA) model was generated from each condition versus fixation contrast maps.

Psychophysiological interaction (PPI) analysis (Friston et al., 1997) was performed in order to assess the context-related differences in functional connectivity between the amygdala and other brain regions, especially the nucleus accumbens. PPI analysis estimates functional connectivity of a pre-defined seed region with the other brain areas by modeling three variables: 1) the task design (psychological variable), 2) task-independent timeseries correlation (physiological variable), and 3) the interaction between these two variables (Friston et al., 1997). This psychophysiological interaction can then be used to calculate functional connectivity estimates between a seed region and other areas of the brain that are dependent on task conditions. Amygdala voxels that were found to show increased activity to negatively modulated surprise vs. positively modulated surprise (GLM_2_) were used as a seed region of interest (ROI) for the PPI analysis. Average timeseries data from these amygdala voxels were deconvolved from the hemodynamic function, multiplied by the onsets of negatively modulated surprise face trials and positively modulated surprise face trials, and then convolved again with the hemodynamic function. This regressor, which represents the interaction between the psychological and physiological signal estimates, was entered into a general linear model, as well as the onsets of each task condition, timeseries data for the seed ROI, and the covariates of no interest. Linear contrast map of negatively modulated surprise versus positively modulated surprise was constructed for each participant, and then submitted to a random effects model.

Given our *a priori* focus on the amygdala, we imposed a corrected significance threshold of *p* < 0.05 over the bilateral amygdala, defined using the Automated Anatomical Labeling atlas (Maldjian et al., 2003). The corrected threshold was determined by Monte Carlo simulations (*n* = 10,000) via *3dClustSim* within AFNI software (Cox, 1996). *3dClustSim*, in conjunction with *3dFWHMx*, allows for a more accurate calculation of corrected thresholds by incorporating the use of advanced spatial autocorrelation function to estimate noise smoothness. The corrected *p* < 0.05 threshold within the amygdala corresponded to uncorrected *p* < 0.005, *k* ≥ 4 voxels (108 mm^3^). In addition, for the PPI analysis, we used a corrected threshold of *p* < 0.05 over the volume of the bilateral NAcc, defined using the Harvard-Oxford probabilistic atlas (Desikan et al., 2006). Using the same procedures described above, corrected *p* < 0.05 was achieved by using uncorrected *p* < 0.005, *k* ≥ 4 voxels (108 mm^3^) for the NAcc. For the all other brain regions, whole-brain corrected *p* < 0.05 threshold was achieved by using uncorrected *p* < 0.001, *k* ≥ 53 voxels (1,431 mm^3^) computed through Monte Carlo simulations.

### Post-Scan Assessment

Upon exiting the scanner, in order to measure each participant’s valence bias, they viewed all 36 individual surprised faces (200 ms per face) sequentially in a randomly shuffled order and asked to rate the valence of each face on a 9-point Likert scale that ranges from -4 to +4 (−4 = *very negative*, +4 = *very positive*). Valence bias refers to a trait-like tendency to view emotionally ambiguous stimuli, such as surprised faces, as positive or negative (Neta, Norris, & Whalen, 2009; Petro et al., 2018). Valence bias was operationally defined as the ratio of negative ratings (< 0) across all 36 faces, in order to facilitate comparisons with previous studies. As a manipulation check, the participants also viewed all 12 sentences (2 s per sentence) sequentially in a randomly shuffled order and were asked to rate the valence on each sentence using the same scale described above. As a final step, participants filled out the State-Trait Anxiety Inventory Form-Y (STAI; Spielberger, Gorsuch, & Lushene, 1988) and the Positive and Negative Affective Schedule (PANAS; Watson, Clark, & Tellegen, 1988). All of the post-scan measurements were used to quantify the individual difference factors that might possibly influence the perceived valence of surprised faces (i.e., valence bias, anxiety, mood). State anxiety scores (STAI-S) were used as a proxy for each participants’ current anxiety levels. Both PANAS negative and positive mood scores were used as an index of current mood. Statistical analyses were performed using SPSS 21 (IBM Corp., Armonk, NY, USA). Data plots were visualized using the Seaborn library (https://doi.org/10.5281/zenodo.3767070) in Python 3.7.1.

## Results

### Behavioral Results

Valence ratings of the surprised faces summarized by six conditions, averaged across all 24 participants were as follows (Figure 1): positive feature + positive context (*M* ± *SD* = 2.1 ± 0.66), positive feature + negative context (−0.85 ± 0.91), ambiguous feature + positive context (1.15 ± 0.9), ambiguous feature + negative context (−2.12 ± 0.74), negative feature + positive context (0.44 ± 1.01), negative feature + negative context (−2.21 ± 0.61). Repeated measures ANOVA results showed a significant main effect of feature (*F*_(2,46)_ = 62.8, *p* < 0.000001, partial *η* = 0.73), a significant main effect of context (*F*_(1,23)_ = 122.1, *p* < 0.000001, partial *η* = 0.84), and a significant interaction (*F*_(2,46)_ = 15.1, *p* < 0.000001, partial *η* = 0.4) of valence ratings, indicating the participants did not wholly depend on contextual cues and utilized subtle facial features during valence computation (Figure 2). The interaction was primarily driven by negative context having a stronger effect on the ambiguous faces. This suggests that contextual information – negative context, in particular – is more important for making valence judgment on ambiguous faces. As ambiguous surprised faces, by definition, are characterized with features that do not offer clear valence signals (Kim et al., 2017), participants would more readily rely on available contextual cues to guide their valence decisions. *Post hoc* analysis revealed that all condition pairs except [ambiguous feature + negative context & negative feature + negative context] were significantly different from each other (Bonferroni corrected *p* < 0.05).

**Figure 2.**
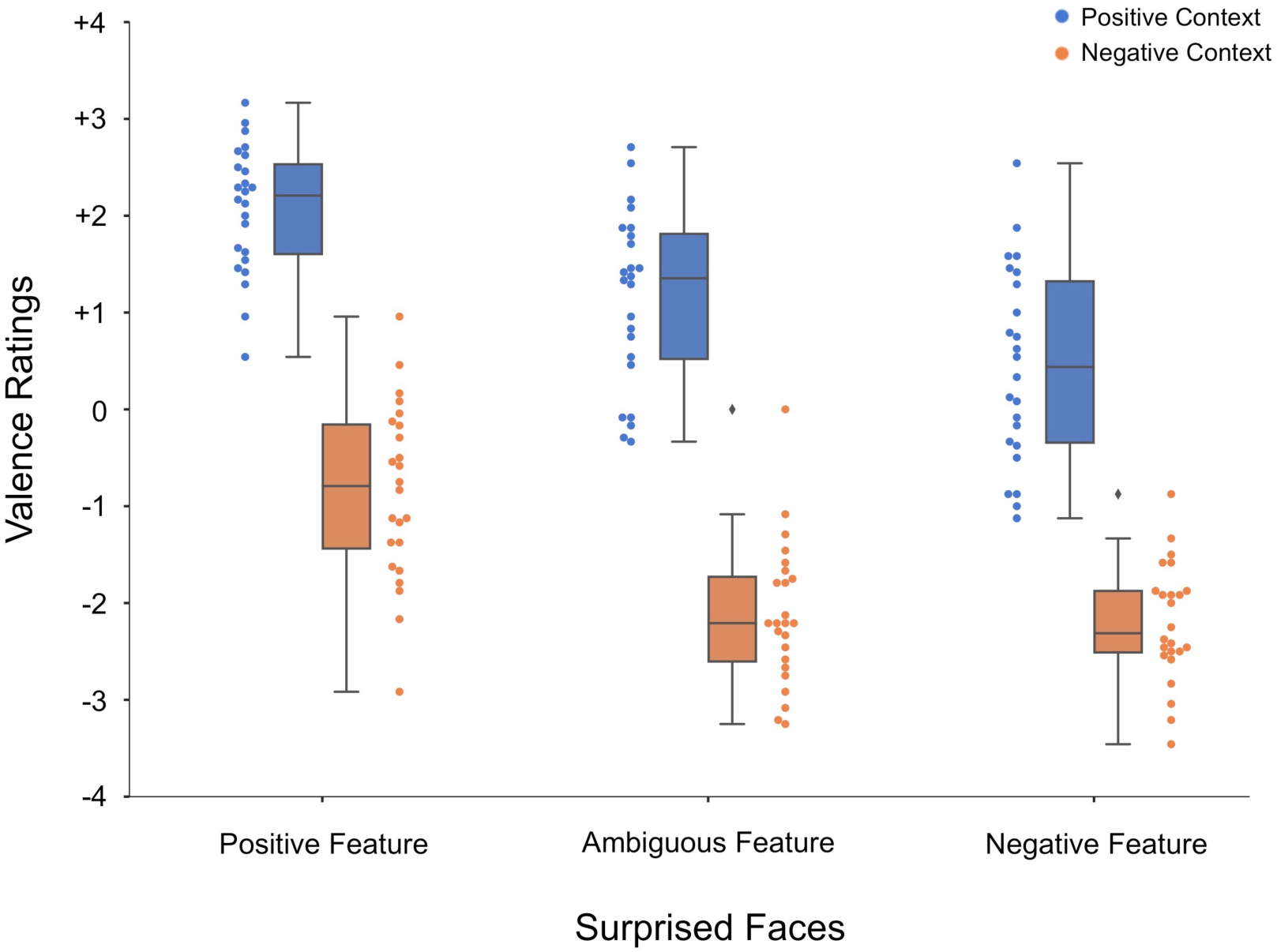
Valence ratings for surprised faces summarized by context and facial features. The box-and-whisker and swarm plots are used to illustrate the clearly distinguishable patterns between positively modulated (blue) and negatively modulated surprised faces (orange), which indicate a substantial main effect of context in shifting the perception of valence in surprise. Main effects of feature as well as the interaction were also significant. Horizontal line within the box represents the mean, the upper- and lower-bounds of the box corresponds to the quartiles of the data, and the whiskers show the rest of the distribution.

Post-scan valence ratings for the faces (without sentences) and sentences (without faces) served as a manipulation check. Consistent with the feature-based labels, the participants rated faces with positive features (0.86 ± 0.87), ambiguous features (−0.69 ± 0.8), or negative features (−1.35 ± 0.57) accordingly (repeated measures ANOVA: *F*_(2,46)_ = 81.5, *p* < 0.000001, partial *η* = 0.78; all *post hoc* pairwise differences were significant at Bonferroni corrected *p* < 0.05). Also, as expected, valence ratings for positive sentences (2.96 ± 0.38) and negative sentences (−2.95 ± 0.45) were significantly different (*t*_(23)_ = 36.4, *p* < 0.000001, *d* = 7.43). Valence bias, quantified as the ratio of negative ratings across 36 surprised faces presented without contextual cues, yielded an average of 0.55 (± 0.14; range 0.36-0.92). Descriptive statistics of the self-report measures of anxiety and mood are as follows: STAI-S (34 ± 9.92), PANAS-P (26.21 ± 5.38), PANAS-N (13.75 ± 5.5).

### fMRI Results

Participant’s valence rating to each face-context pair significantly modulated the activity of the right amygdala (MNI 21, 0, -21; *t*_(23)_ = 3.52, mean *β* = 0.15, *k* = 10 voxels, *d* = 0.76; see Figure 3), indicating that trials with more negative ratings were associated with increased amygdala response. Here, *β* represents the amount of increase in activity per unit of the valence scale. All other brain regions showing significant parametric modulation effects are summarized in Table 1.

**Table 1.** Brain regions showing parametrically modulated BOLD signal response to valence ratings of surprised faces from exploratory whole brain analysis.

| Brain Region | Side | *t* | *x* | *y* | *z* |
| --- | --- | --- | --- | --- | --- |
| *Increased Activity to Negative Ratings* |  |  |  |  |  |
| Lingual Gyrus | L | 6.2 | -18 | -96 | -12 |
| Superior Parietal Lobule | R | 5.5 | 27 | -63 | 57 |
| Postcentral Gyrus | R | 4.7 | 51 | -15 | 54 |
| *Increased Activity to Positive Ratings* |  |  |  |  |  |
| *No brain regions were observed* |  |  |  |  |  |

**Figure 3.**
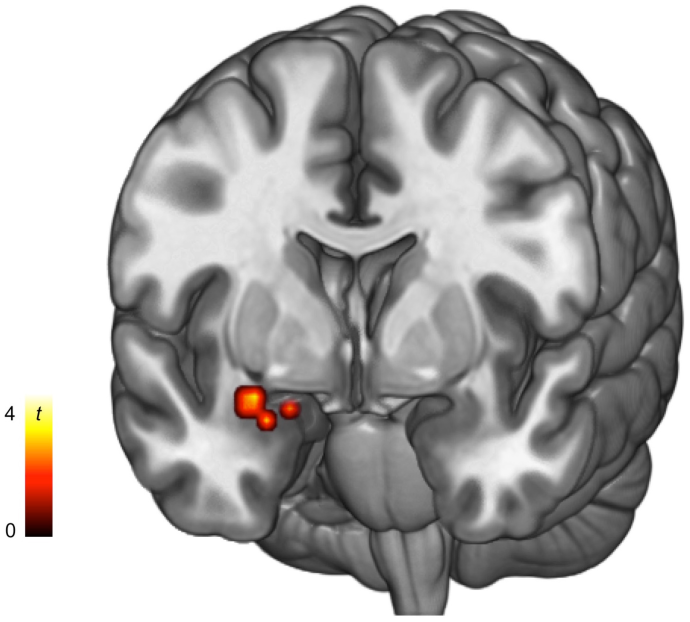
Parametric modulation analysis showing right amygdala activity was increased to trials where the participants responded with more negative ratings.

Consistent with the results from the parametric modulation analysis, a significant increase in right amygdala activity was related to a contrast between negatively modulated surprised faces vs. positively modulated surprised faces (MNI 24, 3, -21; *t*_(23)_ = 3.75, *k* = 20 voxels, *d* = 0.79; Figure 4A). This effect was robust across surprised faces with positive, ambiguous, and negative features. This amygdala voxel cluster showed overlap in 6 voxels with those identified from the parametric modulation analysis. Peak voxels from both clusters were primarily located in the basolateral subnuclei of the amygdala (Tyszka & Pauli, 2016). Not surprisingly, the whole brain activation patterns shown in this contrast, which included the right postcentral gyrus and the right superior parietal lobule, were similar to those found in the parametric modulation analysis, as negatively modulated surprised face trials tend to be when the participants would respond with more negative valence ratings. Neither the surprised faces with negative features vs. positive features contrast (collapsing across negative and positive context to test for feature-based valence effects), nor the negative context vs. positive context contrast (comparing the sentence components) yielded significant brain activity, including the amygdala. No clusters within the amygdala displayed increased activity to the positive vs. negative contrasts.

**Figure 4.**
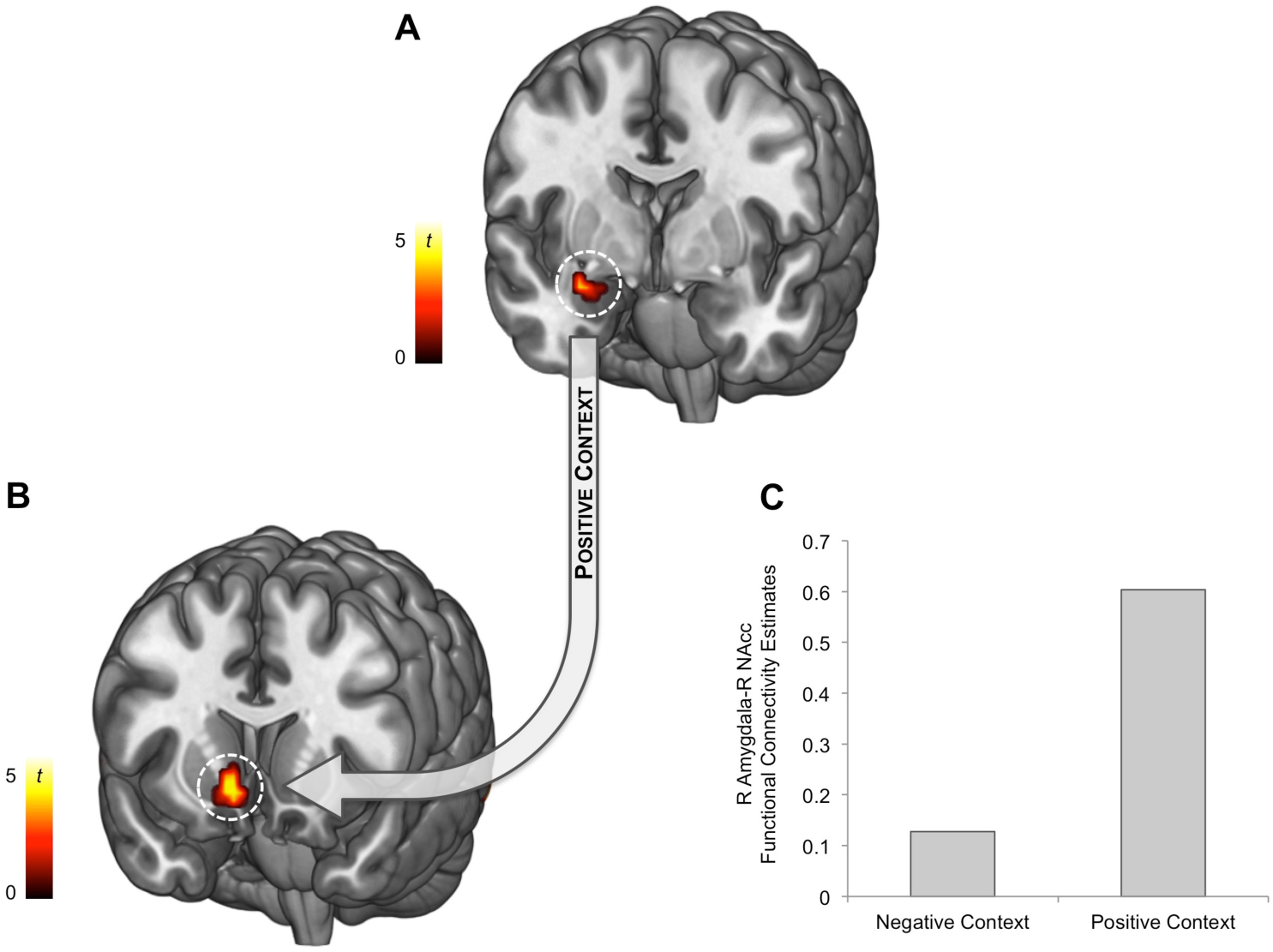
Psychophysiological interaction analysis showing that the functional connectivity of the (A) amygdala and the (B) NAcc was dependent on the valence of the contextual information. (C) Amygdala and the NAcc were functional coupled when processing faces modulated by positive, but not negative context. We note that the bar graphs presented here are for illustration purposes only.

PPI analysis showed that the functional connectivity between the amygdala voxels from GLM_2_ (negatively modulated surprised faces vs. positively modulated surprised faces contrast) with the right ventral striatum was stronger when the faces were paired with positive sentences compared to negative sentences (MNI 9, 15, -6; *t*_(23)_ = 4.16, *k* = 13 voxels, *d* = 0.82; Figure 4B). No other region was differentially coupled with the amygdala during the tasks. Neither the right amygdala activity nor the degree of the right amygdala-NAcc functional connectivity was correlated with valence bias, or self-reported levels of anxiety or mood.

For completeness, the results from the whole brain voxelwise ANOVA are reported in Table 2. Interestingly, *post hoc* examination of brain regions displaying significant interactions between contextual cues and facial features, including the dmPFC (MNI -6, 27, 54; *F*_(2,46)_ = 28.2, *k* = 201 voxels, partial *η* = 0.58; see Figure 5), all showed that this effect was driven by greater response to positive surprised faces that were being modulated by negative sentences compared to positive sentences. This interaction, characterized by robust BOLD response to mismatching information in the dmPFC, is typically observed during cognitive tasks that involve conflict detection (Botvinick, Cohen, & Carter, 2004; Etkin et al., 2006).

**Table 2.** Brain regions showing significant BOLD signal response to the main effects and interaction of the context × feature voxelwise ANOVA from exploratory whole brain analysis.

| Brain Region | Side | *F* | *x* | *y* | *z* |
| --- | --- | --- | --- | --- | --- |
| *Main Effect of Context* |  |  |  |  |  |
| Middle Occipital Gyrus | L | 29.4 | -21 | -102 | -39 |
| Middle Occipital Gyrus | R | 21.0 | 48 | -72 | -9 |
| Postcentral Gyrus | R | 54.3 | 54 | -15 | 54 |
| Superior Parietal Lobule | R | 25.9 | 27 | -63 | 66 |
| *Main Effect of Feature* |  |  |  |  |  |
| *No brain regions were observed* |  |  |  |  |  |
| *Context × Feature Interaction* |  |  |  |  |  |
| Inferior Frontal Gyrus | R | 35.2 | 60 | 21 | 9 |
| Inferior Frontal Gyrus | L | 19.3 | -51 | 18 | 9 |
| Inferior Parietal Lobule | R | 24.9 | 57 | -57 | 54 |
| Dorsomedial Prefrontal Cortex | L | 28.2 | -6 | 27 | 54 |
| Middle Frontal Gyrus | R | 21.5 | 45 | 18 | 42 |

**Figure 5.**
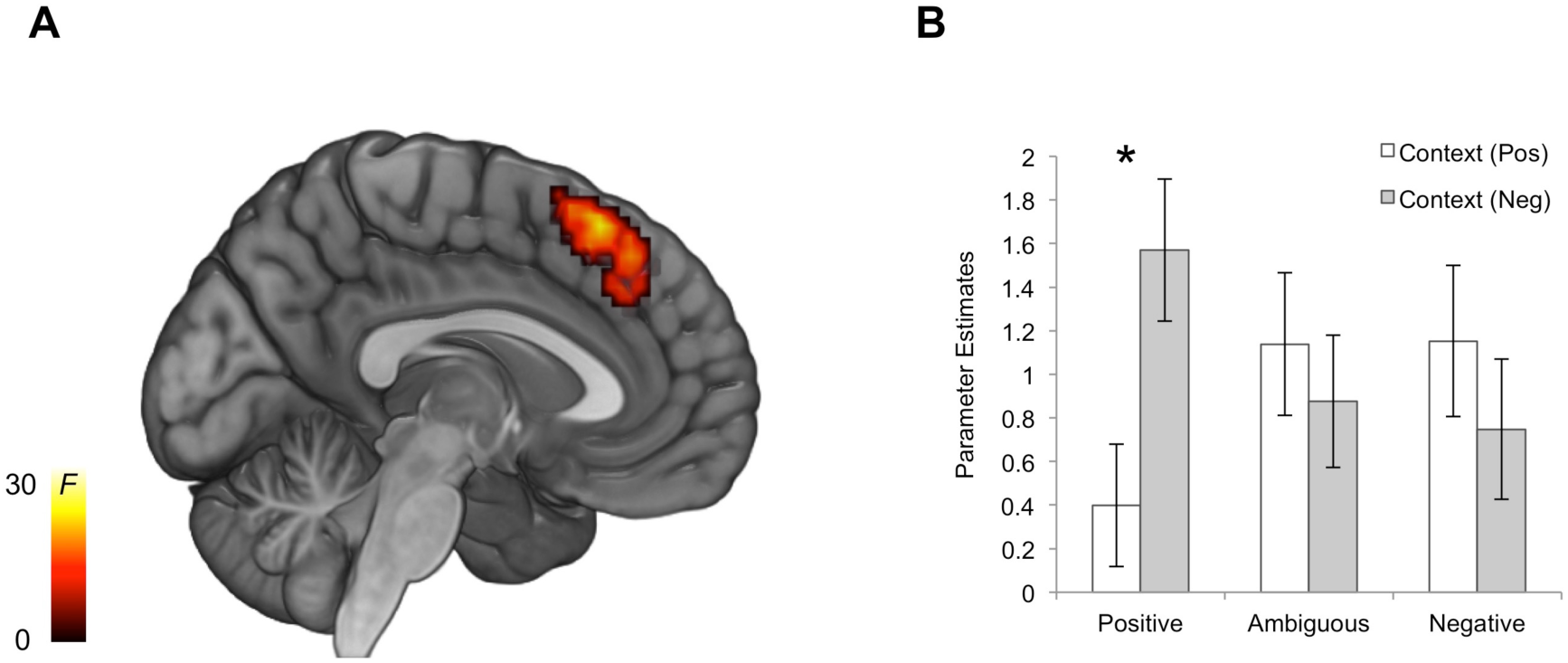
(A) Activity of the dmPFC reflects an interaction between facial features and contextual cues. (B) *Post hoc* analysis showed that this interaction was driven by a significant difference in dmPFC BOLD signal response to positive surprised faces modulated by negative context versus positive context (\**p* < 0.05, Bonferroni corrected). Similar patterns emerged for all other brain regions that showed a significant context × feature interaction effect.

In addition, to examine the specific influence of context on subsequent valence ratings, additional parametric modulation analyses were conducted with the valence rating for each trial as the parametric modulator of hemodynamic responses to the positive and negative sentences, respectively. For positive sentence trials, valence ratings modulated the activity of the right NAcc (MNI 12, 12, -15; *t*_(23)_ = 3.95, mean *β* = 0.17, *k* = 6 voxels, *d* = 0.85), such that trials with more positive ratings were associated with increased NAcc response to the positive sentences. For negative sentence trials, valence ratings modulated the activity of the right postcentral gyrus (MNI 51, -9, 30; *t*_(23)_ = 4.5, mean *β* = 0.28, *k* = 54 voxels, *d* = 0.88), such that trials with more negative ratings were associated with decreased postcentral gyrus response to the negative sentences.

Finally, as both of our amygdala findings were found in the right hemisphere, we sought to test for a potential laterality effect. *Post hoc* tests were performed by creating a left hemisphere version of the two right amygdala findings by flipping the sign of the *x*-coordinates. Then, parameter estimates from the left amygdala voxels were extracted and submitted to paired-sample *t*-tests against those of the right amygdala voxels. In both cases, right amygdala activity was greater than the left (parametric modulation effect: *t*_(23)_ = 2.48, *p* = 0.021, *d* = 0.51; contextual modulation effect: *t*_(23)_ = 2.46, *p* = 0.022; *d* = 0.5).

## Discussion

The present study leveraged two unique characteristics of surprised facial expressions that we observed from our previous research: 1) that their perceived valence is malleable and can be shifted across bipolar valence (positive to negative) while maintaining a consistent level of arousal (Mattek, Wolford, & Whalen, 2017), and 2) that there are subtle, quantifiable features embedded within the face *per se* that guides our perception of valence (Kim et al., 2017). We demonstrated that the valence of surprised facial expressions, which were computed by integrating two sources of information – contextual sentences and facial features – is reflected in the activity of the right amygdala. Importantly, amygdala activity was not found for feature-based signals in the faces or the sentences *per se*. Furthermore, this region of the amygdala was functionally coupled with the NAcc, only when the perceived valence values were shifted due to positive contextual cues. The observed effects were not explained by individual differences in valence bias, mood, or anxiety levels. These data suggest that valence-related amygdala activity resembles the final, subjective judgment of valence, rather than the valence values of each individual component. It also highlights a possible neural mechanism for affective coloring that may be specific for positive valence, via an amygdalostriatal pathway.

First, we replicated the findings from a previous study (Kim et al., 2004) by showing increased amygdala activity in response to surprised faces paired with negative sentences versus positive sentences. Then, we expanded upon these findings in three ways: 1) connecting trial-by-trial behavioral responses with fMRI data using parametric modulation analysis, 2) assessing the amygdalostriatal functional connectivity using PPI analysis, and 3) examining the potential behavioral and neural effects of low-level facial features. As expected, amygdala activity was parametrically modulated by the valence ratings, such that the more negative ratings were associated with greater amygdala BOLD signals. The outcomes from the parametric modulation analysis in particular offer a useful hint about the information that is being held in valence-related amygdala activity. That is, as these amygdala responses are associated with behavioral responses during assigning valence values, they may be reflecting the perceived valence that has already been integrated from available sources of information – namely, contextual cues and facial features. In other words, amygdala activity may be representing the subjective evaluation/judgment regarding valence values, which would be consistent with reports that a subset of amygdala neurons (Wang et al., 2014) or amygdala BOLD activity (Pessoa et al., 2006) follow the subjective judgment of perceived emotional faces, rather than their low-level stimulus properties. This assertion is corroborated by the observation that amygdala activity was not associated with the valence values of contextual cues (i.e., no amygdala responses to negative vs. positive context) or facial features (i.e., no amygdala responses to surprise faces with negative features versus positive features) separately. Considering the findings of our previous fMRI study that did show significant increases in amygdala activity to surprise faces with negative features versus positive features (Kim et al., 2017), the present data demonstrates that valence-related amygdala activity is unlikely to reflect the valence values generated from the bottom-up (or any single source, when multiple sources of information is available); rather, it is likely to represent the final, integrated valence values on which the perceiver has decided. To put it another way, the simplest explanation that reconciles both findings is that the amygdala is representing the final, subjective judgment of valence, be it derived solely from facial features (i.e., Kim et al., 2017) or facial features integrated with contextual cues.

Notably, this area of the amygdala was functionally coupled with the NAcc only during trials with faces paired with positive context, but not negative context. This is consistent with the known role of the NAcc in reward processing and positive affect from both animal research (Berridge & Kringelbach, 2015) and human neuroimaging studies (Wang, Smith, & Delgado, 2016), including our data showing that NAcc responses to positive sentences were increased in trials with more positive ratings. The present findings fit well with evidence from animal work highlighting the importance of amygdala-NAcc interactions via amygdalostriatal pathways in processing positive valence (Beyeler et al., 2018; Namburi et al., 2015; Stuber et al., 2011). One possible interpretation is that in order to perceive a surprised face in a positive light, a more concerted effort between the amygdala and NAcc may be needed to override the primacy of its negative interpretation (Neta & Whalen, 2010; Petro et al., 2018). We note that as PPI analyses cannot imply directionality on its own, interpretations of functional connectivity should be made with caution.

While we have focused on the amygdala’s role in managing valence information for the purposes of the current study, we are of course not suggesting that the amygdala is the sole neural site for such processes. Rather, it is becoming increasingly important to consider the role of other brain regions for encoding valence. In this regard, it would be useful to compare the present study with a number of notable works on the neural bases for valence encoding. An fMRI study by Jin and colleagues (2015) used odor stimuli that ranged from very pleasant (e.g., lemon) to very unpleasant (e.g., sulfurous), and found that the activity of the amygdala represented a unidimensional valence code. Thus, the valence reported by the participants is almost exclusively based on strong, visceral reactions (Jin et al., 2015). Skerry and Saxe (2014), on the other hand, used a different approach to examine the neural representation of valence. They focused on how we infer emotions from situations and context (e.g., short clips of animations depicting a scenario where the character’s emotional state is affected), and found that the activity of the dmPFC represented the high-level conceptual representation of valence (Skerry & Saxe, 2014). Due to the involvement of top-down inferential process, the valence being studied in the latter study may be more comparable to high-level social cognition such as theory of mind, rather than the bottom-up, stimulus-driven approach used by the former. These two studies and others (e.g., Comte et al., 2016; McRae et al., 2012; Ochsner et al., 2009) illustrate the importance of considering *how* the valence was generated in exploring its neural underpinnings. The current study is situated in between, as we examined the neural correlates of a valence computation process that requires both bottom-up generated reactions and top-down inference.

Exploratory analyses of the whole brain yielded responses of the inferior frontal gyrus, inferior parietal lobule, middle frontal gyrus, and dmPFC during the processing of conflicting information between the facial features and contextual cues. We are careful not to overly interpret these *post hoc* findings, but it is notable that the dmPFC offers an interesting explanation in particular, as it and the adjacent dorsal anterior cingulate cortex are responsive to conflicting information more generally (Botvinick, Cohen, & Carter, 2004; Etkin et al., 2006). More recently, the dmPFC and the dorsal anterior cingulate cortex have been suggested to be broadly involved in performing tasks and detecting/selecting salient stimuli, as a part of the task activation network (Dosenbach et al., 2006) and the salience network (Seeley et al., 2007), respectively. In light of this view of the dmPFC, it seems plausible to understand that positive stimulus features within a negative context may provide a mismatch in information, even if the features are subtle, and likely requires a greater deliberate effort to change the valence. Moreover, since the negative context was followed by positive faces, the ensuing unexpected positive surprise could be viewed as positive prediction error. This interpretation is consistent with studies showing that the dmPFC and the dorsal anterior cingulate cortex are able to encode positive prediction errors (Behrens et al., 2008; Hayden et al., 2011). Putting the findings from the present study together, the following speculative suggestion can be made: during valence computation, whether the amygdala communicates with the NAcc depends on the available contextual cue, which is integrated with the low-level affective information from the features to produce the final valence value. Throughout this process, the dmPFC may be particularly sensitive to conflicting information from the two sources, and potentially acts to resolve this mismatch.

### Limitations

Of course, the present study is not without limitations that could be addressed in future studies. An alternative reason for the lack of feature-based valence effects in the amygdala, despite the significant main effect of facial features on a behavioral level, may be attributable to the experimental design used in the present study. Participants were explicitly evaluating the valence of the faces, and it has been suggested that an explicit emotional task may have substantial mitigating impact on amygdala activity (Costafreda et al., 2008). In contrast, our previous fMRI experiment involved simple passive viewing of faces, which may have allowed us to observe feature-based valence effects in the amygdala (Kim et al., 2017). Future studies employing an orthogonal task may be able to locate the fine-grained valence information based on subtle facial features encoded in the brain, amidst the substantial effect of context. In addition, yet another possibility is that it may not be feasible to observe brain activity that encodes each individual component of valence computation (e.g., valence value of faces *per se*), due to the limited temporal resolution of fMRI and lagged response of BOLD signals. If valence computation requires valence values for each individual component as its input, it follows then that this information must be encoded in the brain. One way to locate this valence code in future studies is the inclusion of trials without contextual cues, such that feature-based valence effect within the amygdala and other brain regions could be estimated.

Another potential issue is that the subjective valence ratings of the contextually modulated faces generally follow the clear valence of the sentences. Future investigations may employ more subtle contextual cues in order to better delineate the valence of the individual components from the integrated valence. The unexpected right-lateralized effect of the amygdala observed in the present data does not lend itself to a straightforward interpretation. If anything, the inclusion of language in the form of contextual cues would have been likely associated with the left amygdala (Costafreda et al., 2008). Future investigations with larger sample size and systematical manipulation of study components (e.g., verbal vs. non-verbal context, rating task vs. passive viewing) could address the possibility of hemispheric laterality in the amygdala. Also, due to the purposes of the present study, we chose to primarily focus on the amygdala’s role in encoding valence. As increasing evidence points to the likelihood that valence information may be represented as distributed brain activity patterns (Chang et al., 2015; Chikazoe et al., 2014; Kim, Shinkareva, & Wedell, 2017; Knutson, Katovich, & Suri, 2014; Lindquist et al., 2016; Miskovic & Anderson 2018), the idea of a valence code possibly being represented in a distributed manner should be considered. Future work would be able to build on the present findings and extend the scope to assess whole brain voxels using an MVPA approach. Finally, our recent fMRI work suggests different brain regions that seemingly encode valence may emerge depending on how core affective dimensions are defined (e.g., valence/arousal dimensions vs. positivity/negativity dimensions) (Mattek et al., 2020). One suggested approach is to consider all four affective dimensions when collecting and analyzing data to further elucidate the link between valence values and brain activity.

### Conclusions

To summarize, the current study showed that, when given multiple sources of information, our ability to incorporate and integrate them to produce a subjective assessment of perceived valence as reflected in rating behavior is also represented in the amygdala. Findings from the present study imply that the amygdala isn’t simply encoding valence from a single source, but is able to update and modify by incorporating contextual cues. Integrating positive contextual cues in particular involves functional coupling of the amygdala and the NAcc. The present data shed light on the role of the amygdala in valence computation, which in turn offers a possible neural mechanism that may underpin how affective information from multiple sources are integrated and processed to guide our behavior.

## Acknowledgements

We thank Randi Bennett and Kimberly Solomon for their assistance with data collection and organization. The present study is a part of Justin Kim’s doctoral dissertation work conducted at Dartmouth College. This work was supported by the National Institute of Mental Health under Grant F31MH090672 and R01MH080716.

## Open Practices Statement

None of the data or materials for the experiments reported here is publicly available due to lack of informed consent and ethical approval, but is available on request by qualified scientists. None of the experiments was preregistered.

## References

Alheid, G. F., & Heimer, L. (1988). New perspectives in basal forebrain organization of special relevance for neuropsychiatric disorders: the stratopallidal, amygdaloid, and corticopetal components of substantia innominate. Neuroscience, 27, 1–39. https://doi.org/10.1016/0306-4522(88)90217-5

Anders, S., Eippert, F., Weiskopf, N., & Veit, R. (2008). The human amygdala is sensitive to valence of pictures and sounds irrespective of arousal: an fMRI study. Social Cognitive and Affective Neuroscience, 3, 233–243. https://doi.org/10.1093/scan/nsn017

Behrens, T. E. J., Hunt, L. T., Woolrich, M. W., & Rushworth, M. F. S. (2008). Associative learning of social value. Nature, 456, 245–249. https://doi.org/10.1038/nature07538

Belova, M. A., Paton, J. J., & Salzman, C. D. (2008). Moment-to-moment tracking of state value in the amygdala. Journal of Neuroscience, 28, 10023–10030. https://doi.org/10.1523/JNEUROSCI.1400-08.2008

Berridge, K. C., & Kringelbach, M. L. (2015). Pleasure systems in the brain. Neuron, 86, 646–664. https://doi.org/10.1016/j.neuron.2015.02.018

Beyeler, A., Chang, C. J., Silvestre, M., Lévêque, C., Namburi, P., Wildes, C. P., … & Tye, K. M. (2018). Organization of valence-encoding and projection-defined neurons in the basolateral amygdala. Cell Reports, 22, 905–918. https://doi.org/10.1016/j.celrep.2017.12.097

Botvinick, M. M., Cohen, J. D., Carter, C. S. (2004). Conflict monitoring and anterior cingulate cortex: an update. Trends in Cognitive Sciences, 2004, 539–546. https://doi.org/10.1016/j.tics.2004.10.003

Cacioppo, J. T., Petty, R. E., Losch, M. E., & Kim, H. S. (1986). Electromyographic activity over facial muscle regions can differentiate the valence and intensity of affective reactions. Journal of Personality and Social Psychology, 50, 260–268. https://doi.org/10.1037/0022-3514.50.2.260

Chang, L. J., Gianaros, P. J., Manuck, S. B., Krishnan, A., & Wager, T. D. (2015). A sensitive and specific neural signature for picture-induced negative affect. PLoS Biology, 13, e1002180. https://doi.org/10.1371/journal.pbio.1002180

Chikazoe, J., Lee, D. H., Kriegeskorte, N., & Anderson, A. K. (2014). Population coding of affect across stimuli, modalities and individuals. Nature Neuroscience, 17, 1114–1122. https://doi.org/10.1038/nn.3749

Clithero, J. A., & Rangel, A. (2013). Informatic parcellation of the network involved in the computation of subjective value. Social Cognitive and Affective Neuroscience, 9, 1289–1302. https://doi.org/10.1093/scan/nst106

Comte, M., Schön, D., Coull, J. T., Reynaud, E., Khalfa, S., Belzeaux, R., … & Fakra, E. (2016). Dissociating bottom-up and top-down mechanisms in the cortico-limbic system during emotion processing. Cerebral Cortex, 26, 144–155. https://doi.org/10.1093/cercor/bhu185

Costafreda, S. G., Brammer, M. J., David, A. S., & Fu, C. H. Y. (2008). Predictors of amygdala activation during the processing of emotional stimuli: A meta-analysis of 385 PET and fMRI studies. Brain Research Reviews, 58, 57–70. https://doi.org/10.1016/j.brainresrev.2007.10.012

Cox R. W. (1996). AFNI: software for analysis and visualization of functional magnetic resonance neuroimages. Computational Biomedical Research, 29, 162–173. https://doi.org/10.1006/cbmr.1996.0014

Cunningham, W. A., Van Bavel, J. J., & Johnsen, I. R. (2008). Affective flexibility: evaluative processing goals shape amygdala activity. Psychological Science, 19, 152–160. https://doi.org/10.1111/j.1467-9280.2008.02061.x

Desikan, R. S., Ségonne, F., Fischl, B., Quinn, B. T., Dickerson, B. C., Blacker, D., … & Killiany, R. J. (2006). An automated labeling system for subdividing the human cerebral cortex on MRI scans into gyral based regions of interest. Neuroimage, 31, 968–980. https://doi.org/10.1016/j.neuroimage.2006.01.021

Dosenbach, N. U. F., Visscher, K. M., Palmer, E. D., Miezin, F. M., Wenger, K. K., Kang, H. C., … & Petersen, S. E. (2006). A core system for the implementation of task sets. Neuron, 50, 799–812. https://doi.org/10.1016/j.neuron.2006.04.031

Ekman, P. F., & Friesen, W. V. (1976). Pictures of facial affect. Palo Alto: Consulting Psychologists Press.

Etkin, A., Egner, T., & Kalisch, R. (2011). Emotional processing in anterior cingulate and medial prefrontal cortex. Trends in Cognitive Sciences, 15, 82–83. https://doi.org/10.1016/j.tics.2010.11.004

Etkin, A., Egner, T., Peraza, D. M., Kandel, E. R., & Hirsch, J. (2006). Resolving emotional conflict: A role of the rostral anterior cingulate cortex in modulating activity in the amygdala. Neuron, 51, 871–882. https://doi.org/10.1016/j.neuron.2006.07.029

First, M., Spitzer, M., Williams, J., & Gibbon, M. (1995). Structured clinical interview for DSM-IV (SCID). Washington, DC: American Psychiatric Association.

Friston, K., Buechel, C., Fink, G., Morris, J. Rolls, E., & Dolan, R. (1997). Psychophysiological and modulatory interactions in neuroimaging. Neuroimage, 6, 218–229. https://doi.org/10.1006/nimg.1997.0291

Gottfried, J. A., O’Doherty, J., & Dolan, R. J. (2002). Appetitive and aversive olfactory learning in humans studied using event-related functional magnetic resonance imaging. Journal of Neuroscience, 22, 10829–10837. https://doi.org/10.1523/JNEUROSCI.22-24-10829.2002

Grupe, D. W., Schaefer, S. M., Lapate, R. C., Schoen, A J., Gresham, L. K., Mumford, J. A., … & Davidson, R. J. (2018). Behavioral and neural indices of affective coloring for neutral social stimuli. Social Cognitive and Affective Neuroscience, 13, 310–320. https://doi.org/10.1093/scan/nsy011

Hayden, B. Y., Heilbronner, S. R., Pearson, J. M., & Platt, M. L. (2011). Surprise signals in anterior cingulate cortex: neuronal encoding of unsigned reward prediction errors driving adjustment in behavior. Journal of Neuroscience, 31, 4178–4187. https://doi.org/10.1523/JNEUROSCI.4652-10.2011

Jin, J., Zelano, C., Gottried, J. A., & Mohanty, A. (2015). Human amygdala represents the complete spectrum of subjective valence. Journal of Neuroscience, 35, 15145–15156. https://doi.org/10.1523/JNEUROSCI.2450-15.2015

Kim, H., Somerville, L. H., Johnstone, T., Polis, S., Alexander, A. L., Shin, L. M., & Whalen, P. J. (2004). Contextual modulation of amygdala responsivity to surprised faces. Journal of Cognitive Neuroscience, 16, 1730–1745. https://doi.org/10.1162/0898929042947865

Kim, J., Shinkareva, S. V., Wedell, D. H. (2017). Representations of modality-general valence for videos and music derived from fMRI data. Neuroimage, 148, 42–54. https://doi.org/10.1016/j.neuroimage.2017.01.002

Kim, M. J., Mattek, A. M., Bennett, R. H., Solomon, K. M., Shin J., & Whalen, P. J. (2017). Human amygdala tracks a feature-based valence signal embedded within the facial expressions of surprise. Journal of Neuroscience, 37, 9510–9518. https://doi.org/10.1523/JNEUROSCI.1375-17.2017

Knutson, B., Katovich, K., & Suri, G. (2014). Inferring affect from fMRI data. Trends in Cognitive Sciences, 18, 422–428. https://doi.org/10.1016/j.tics.2014.04.006

Lang, P. J., Bradley, M. M., & Cuthbert, B. N. (1998). Emotion, motivation, and anxiety: Brain mechanisms and psychophysiology. Biological Psychiatry, 44, 1248–1263. https://doi.org/10.1016/s0006-3223(98)00275-3

Lapate, R. C., Rokers, B., Tromp, D. P. M., Orfali, N. S., Oler, J. A., Doran, S. T., … & Davidson, R. J. (2016). Awareness of emotional stimuli determines the behavioral consequences of amygdala activation and amygdala-prefrontal connectivity. Scientific Reports, 6, 25826. https://doi.org/10.1038/srep25826

Lindquist, K. A., Satpute, A. B., Wager, T. D., Weber, J., & Barrett, L. F. (2016). The brain basis of positive and negative affect: evidence from a meta-analysis of the human neuroimaging literature. Cerebral Cortex, 26, 1910–1922. https://doi.org/10.1093/cercor/bhv001

Lundqvist, D., Flykt, A., & Ohman, A. (1998). The Karolinska Directed Emotional Faces-KDEF, CD-ROM from the Department of Clinical Neuroscience, Psychology Section, Karolinska Institutet.

Maldjian, J. A., Laurienti, P. J., Kraft, R. A., & Burdette, J. H. (2003). An automated method for neuroanatomic and cytoarchitectonic atlas-based interrogation of fMRI data sets. Neuroimage, 19, 1233–1239. https://doi.org/10.1016/S1053-8119(03)00169-1

Mattek, A. M., Burr, D. A., Shin, J., Whicker, C. L., & Kim, M. J. (2020). Identifying the representational structure of affect using fMRI. Affective Science. https://doi.org/10.1007/s42761-020-00007-9

Mattek, A. M., Wolford, G. L., & Whalen, P. J. (2017). A mathematical model captures the structure of subjective affect. Perspectives in Psychological Science, 12, 508–526. https://doi.org/10.1177/1745691616685863

McRae, K., Misra, S., Prasad, A. K., Pereira, S. C., & Gross, J. J. (2012). Bottom-up and top-down emotion generation: implications for emotion regulation. Social Cognitive and Affective Neuroscience, 7, 253–262. https://doi.org/10.1093/scan/nsq103

Miskovic, V., & Anderson, A. K. (2018). Modality general and modality specific coding of hedonic valence. Current Opinion in Behavioral Sciences, 19, 91–97. https://doi.org/10.1016/j.cobeha.2017.12.012

Morrison, S. E., & Salzman, C. D. (2009). The convergence of information about rewarding and aversive stimuli in single neurons. Journal of Neuroscience, 29, 11481–11483. https://doi.org/10.1523/JNEUROSCI.1815-09.2009

Namburi, P., Beyeler, A., Yorozu, S., Calhoon, G. G., Halbert, S. A., Wichmann, R., … & Tye, K. M. (2015). A circuit mechanism for differentiating positive and negative associations. Nature, 520, 675–678. https://doi.org/10.1038/nature14366

Neta, M., Norris, C. J., & Whalen, P. J. (2009). Corrugator muscle responses to surprised facial expressions are associated with individual differences in positivity-negativity bias. Emotion, 9, 640–648. https://doi.org/10.1037/a0016819

Neta, M., & Whalen, P. J. (2010). The primacy of negative interpretations when resolving the valence of ambiguous facial expressions. Psychological Science, 21, 901–907. https://doi.org/10.1177/0956797610373934

Ochsner, K. N., Ray, R. R., Hughes, B., McRae, K., Cooper, J. C., Weber, J., … & Gross, J. J. (2009). Bottom-up and top-down processes in emotion generation: common and distinct neural mechanisms. Psychological Science, 20, 1322–1331. https://doi.org/10.1111/j.1467-9280.2009.02459.x

Paton, J. J., Belova, M. A., Morrison, S. E., & Salzman, C. D. (2006). The primate amygdala represents the positive and negative value of visual stimuli during learning. Nature, 439, 865–870. https://doi.org/10.1038/nature04490

Petro, N. M., Tong, T. T., Henley, D. J., & Neta, M. (2018). Individual differences in valence bias: fMRI evidence of the initial negativity hypothesis. Social Cognitive and Affective Neuroscience, 13, 687–698. https://doi.org/10.1093/scan/nsy049

Pessoa, L., Japee, S., Sturman, D., & Ungerleider, L. G. (2006). Target visibility and visual awareness modulate amygdala responses to fearful faces. Cerebral Cortex, 16, 366–375. https:/doi.org/10.1093/cercor/bhi115

Phelps, E. A., & LeDoux, J. E. (2005). Contributions of the amygdala to emotion processing: from animal models to human behavior. Neuron, 48, 175–187. https://doi.org/10.1016/j.neuron.2005.09.025

Pitkänen, A. (2000). Connectivity of the rat amygdaloid complex. In: J. P. Aggleton (Ed.), The Amygdala: A Functional Analysis, 2nd Edition (pp. 31–116). New York: Offord university Press.

Poldrack, R. A., Baker, C. I., Durnez, J., Gorgolweski, K. J., Matthews, P. M., Munafò, M. R., … & Yarkoni, T. (2017). Scanning the horizon: towards transparent and reproducible neuroimaging research. Nature Reviews Neuroscience, 18, 115–126.

Reynolds, S. M., & Berridge, K. C. (2008). Emotional environments retune the valence of appetitive versus fearful functions in nucleus accumbens. Nature Neuroscience, 11, 423–425. https://doi.org/10.1038/nn2061

Rigoard, P., Buffenoir, K., Jaafari, N., Giot, J. P., Houeto, J. L., Mertens, P., … & Bataille, B. (2011). The accumbofrontal fasciculus in the human brain: a microsurgical anatomical study. Neurosurgery, 68, 1102–1111. https://doi.org/10.1227/NEU.0b013e3182098e48

Russell, J. A. (1980). A circumplex model of affect. Journal of Personality and Social Psychology, 39, 1161–1178. https://doi.org/10.1037/h0077714

Seeley, W. W., Menon, V., Schatzberg, A. F., Keller, J., Glover, G. H., Kenna, H., Reiss, A. K., … & Greicius, M. D. (2007). Dissociable intrinsic connectivity networks for salience processing and executive control. Journal of Neuroscience, 27, 2349–2356. https://doi.org/10.1523/JNEUROSCI.5587-06.2007

Skerry, A. E., & Saxe, R. (2014). A common neural code for perceived and inferred emotion. Journal of Neuroscience, 34, 15997–16008. https://doi.org/10.1523/JNEUROSCI.1676-14.2014

Spielberger, C. D., Gorsuch, R. L., & Lushene, R. E. (1988). STAI-Manual for the State Trait Anxiety Inventory. Palo Alto, CA: Consulting Psychologists Press.

Stuber, G. D., Sparta, D. R., Stamatakis, A. M., van Leeuwen, W. A., Hardjoprajitno, J. E., Cho, S., … & Bonci, A. (2011). Excitatory transmission from the amygdala to nucleus accumbens facilitates reward seeking. Nature, 475, 377–380. https://doi.org/10.1038/nature10194

Tottenham, N., Tanaka, J. W., Leon, A. C., McCarry, T., Nurse, M., Hare, T. A., … & Nelson, C. (2009). The NimStim set of facial expressions: Judgments from untrained research participants. Psychiatry Research, 168, 242–249. https://doi.org/10.1016/j.psychres.2008.05.006

Tyszka, J. M., & Pauli, W. M. (2016). In vivo delineation of subdivisions of the amygdaloid complex in a high-resolution group template. Human Brain Mapping, 37, 3979–3998. https://doi.org/10.1002/hbm.23289

Wang, K. S., Smith, D. V., & Delgado, M. R. (2016). Using fMRI to study reward processing in humans: past, present, and future. Journal of Neurophysiology, 115, 1664–1678. https://doi.org/10.1152/jn.00333.2015

Wang, S., Tudusciuc, O., Mamelak, A. N., Ross, I. B., Adolphs, R., & Rutishauser, U. (2014). Neurons in the human amygdala selective for perceived emotion. Proceedings of the National Academy of Sciences, 111, E3110–3119. https://doi.org/10.1073/pnas.1323342111

Watson, D., Clark, L. A., & Tellegen, A. (1988). Development and validation of brief measures of positive and negative affect: The PANAS scales. Journal of Personality and Social Psychology, 54, 1063–1070. https://doi.org/10.1037/0022-3514.54.6.1063

